# Peripersonal and reaching space differ: evidence from their spatial extent and multisensory facilitation pattern

**DOI:** 10.1101/2020.06.01.127282

**Authors:** A. Zanini, I. Patané, E. Blini, R. Salemme, E. Koun, A. Farnè, C. Brozzoli

## Abstract

Peripersonal space (PPS) is a multisensory representation of the space near body parts facilitating interactions with the close environment. Studies on non-human and human primates converge in showing that PPS is a body-part-centred representation that guides actions. Because of these characteristics, growing confusion conflates peripersonal and arm-reaching space (ARS), that is the space one’s arm can reach. Despite neuroanatomical evidence favors their distinction, no study has contrasted directly their respective extent and behavioral features. Here, in five experiments (N=140) we found that PPS differs from ARS, as evidenced both by participants’ spatial and temporal performance and its modeling. We mapped PPS and ARS using both their respective gold standard tasks and a novel multisensory facilitation paradigm. Results show that 1) PPS is smaller than ARS; 2) multivariate analyses of spatial patterns of multisensory facilitation predict participants’ hand locations within ARS; 3) the multisensory facilitation map shifts isomorphically following hand positions, revealing hand-centred coding of PPS, therefore pointing to a functional similarity to the receptive fields of monkeys’ multisensory neurons. A control experiment further corroborated these results and additionally ruled out the orienting of attention as driving mechanism for the increased multisensory facilitation near the hand. In sharp contrast, ARS mapping results in a larger spatial extent, with undistinguishable patterns across hand positions, cross-validating the conclusion that PPS and ARS are distinct spatial representations. These findings urge for a refinement of theoretical models of PPS, which is relevant to constructs as diverse as self-representation, social interpersonal distance, and motor control.

## Introduction

Seminal studies described multisensory neurons in primates’ fronto-parietal regions coding for the space surrounding the body, called *peripersonal space* (PPS) (Colby et al., 1993; Graziano & Gross, 1993; Rizzolatti et al., 1981a, 1981b). These neurons display visual receptive fields anchored to tactile ones and protruding over a limited area (∼5-30 cm) from specific body-parts (e.g., the hand) (Graziano & Gross, 1993; Rizzolatti et al., 1981a, 1981b). Neuroimaging results in humans are in line with these findings: ventral and anterior intraparietal sulcus, ventral and dorsal premotor cortices and putamen integrate visual, tactile and proprioceptive signals, allowing for a body-part-centred representation of space (Brozzoli et al., 2011, 2012). Behaviorally, visual stimuli modulate responses to touches of the hand more strongly when presented near compared to far from it (Farnè et al., 2005; Làdavas & Farnè, 2004; Serino et al., 2015; Spence et al., 2004), a mechanism proposed to subserve both defensive (de Haan et al., 2016; Graziano & Cooke, 2006) and acquisitive aims (Brozzoli et al., 2009, 2010, 2014; Patané et al., 2019; Vignemont & Iannetti, 2014).

As a multisensory interface guiding interactions with the environment, PPS shares some characteristics with the arm-reaching space (ARS), the space reachable by extending the arm without moving the trunk (Coello et al., 2008). In humans, ARS tasks typically require to judge the reachability of a stimulus (Carello et al., 1989; Coello & Iwanow, 2006). Despite their anatomo-functional differences (Desmurget et al., 1999; Filimon, 2010; Lara et al., 2018; Pitzalis et al., 2013), some research on human PPS grew apart from the original electrophysiological findings and conflated ARS and PPS (Coello et al., 2008; Iachini et al., 2014; Vieira et al., 2020). However, multisensory stimuli within ARS and close to the hand activate neural areas typically associated with PPS, whereas the same stimuli within ARS, but far from the hand, do not (Brozzoli et al., 2012; Graziano et al., 1994). To date, no empirical evidence exists to distinguish these spatial representations. The consequences of this conflation on spatial models of multisensory facilitation have been so far neglected, despite the crucial role it plays for sensorimotor control (Makin et al., 2017; Suminski et al., 2009, 2010) and the study of the bodily self (Blanke et al., 2015; Makin et al., 2008).

Here we leveraged empirical outcomes to disentangle two alternative theoretical models, hypothesizing that PPS and ARS are either identical or distinct spatial representations. To ensure fair comparative bases to this purpose, and to allow making clear alternative predictions, we set two pre-requisites: the first is not to oppose PPS and ARS in the context of different functions; the second is to test both spaces with reference to the same body-part. Thus, in Experiment I we used a tactile detection task and computed multisensory (visuo-tactile) facilitation, a typical proxy of PPS extent. In Experiment II, we used a reachability judgement task and computed the point of subjective equality (PSE), a typical estimate of the ARS extent (Bourgeois & Coello, 2012). As visual and tactile stimuli were harmless and semantically neutral, our tasks were devoid of any defensive or social function. In addition, both PPS and ARS tasks were applied in reference to the hand, as PPS has been shown to be hand-centred (di Pellegrino et al., 1997) and what we can reach (ARS) is defined by how far our hand can get (Coello & Iwanow, 2006), thus fulfilling the criteria for a fair comparison. Two additional experiments manipulated hand vision (visible or not) and position (close or distant), to progressively equate the reachability task to the multisensory conditions of Experiment I.

Following this rationale, if PPS and ARS are equal, we should observe similar spatial extents from multisensory facilitation and reachability estimates. In addition, we should observe facilitation from all visual stimuli falling within ARS independently of hand position. Conversely, we should measure different spatial extents and observe multisensory facilitation only for stimuli near the hand, as a function of its position, resulting in specific and distinguishable spatial patterns of multisensory facilitation.

## Experiment I

### Methods

#### Participants

We calculated our sample size with G*Power 3.1.9.2, setting the 10*2 (V-Position*Hand Position) within-interaction for a RM ANOVA hypothesizing a power of 0.85, an α = 0.05 and a correlation of 0.5 between the measures. Considering a medium-low effect size (0.20, see Holmes, Martin, Mitchell, Noorani, & Thorne, 2020), we needed to recruit at least 23 participants per study. All participants were right-handed, as evaluated via the Edinburgh Handedness Test (mean score 82%). Twenty-seven subjects (13 females; mean age =26.12, range=20-34; mean arm length=79.41 ± 5.83 cm, measured from the acromion to the tip of the right middle finger) participated in Experiment I.

All participants reported normal or corrected-to-normal vision, normal tactile sensitivity and no history of psychiatric disorders. They gave their informed consent before taking part in the study, which was approved by the local ethics committee (Comité d’Evaluation de l’Ethique de l’Inserm, n° 17-425, IRB00003888, IORG0003254, FWA00005831) and was in line with the Declaration of Helsinki. Participants were paid 15 euros.

#### Stimuli and Apparatus

*Visual stimuli* were identical for both the experiments. We used a projector (Panasonic PT-LM1E_C) to present a 2D grey circle (RGB = 32, 32, 32) in one of 10 positions, ranging from near to far from the body. The diameter of the grey circle was corrected for retinal size with the formula:

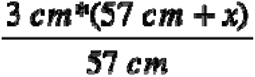

where 3 cm is the diameter of the circle, 57 cm is the distance from the eye at which 1° of the visual field roughly corresponds to 1 cm and *x* is the distance of the centre of the stimulus from the point at 57 cm. Visual stimulus duration was 500 ms. The fixation cross (2.5 cm) was projected along the body’s sagittal axis (see Figure 1). The ten positions were calibrated such that the sixth one corresponded to the objective limit of reachability for each participant. We ensured this before the experiment: participants stayed with eyes closed, their head on a chinrest (30 cm high) and placed their right hand as far as possible on the table. Starting from the sixth position, 4 positions were computed beyond the reachable limit and 5 closer to the participant’s body, 8 cm rightward with respect to the body’s sagittal axis. Positions, uniformly separated by 9 cm, spanned along 90 cm of space and were labelled V-*P1* to V-*P10,* from the closest to the farthest (see Figure 1).

**Figure 1:**
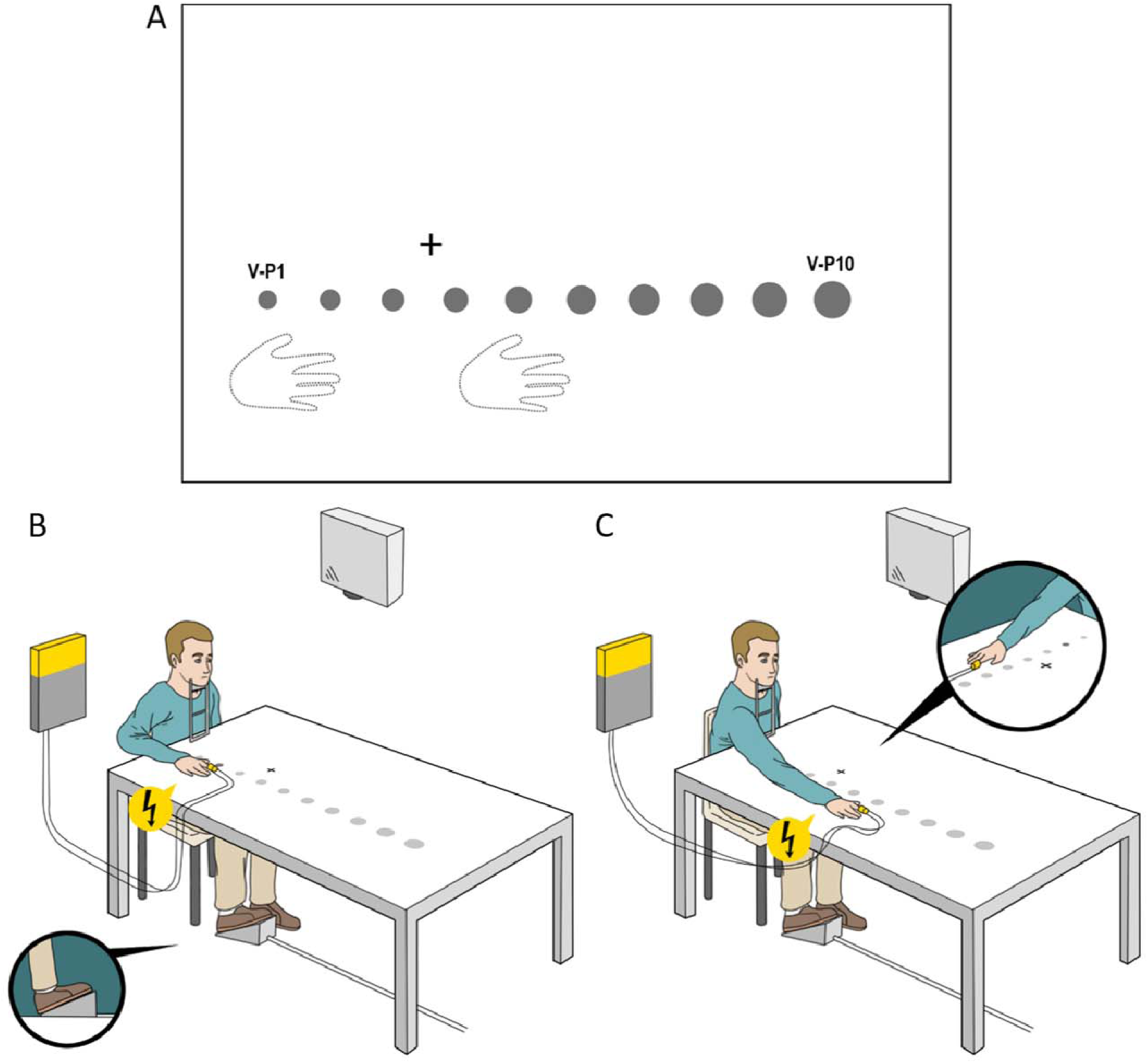
Experimental setup across experiments. <u>a)</u> Positions of right hand, fixation cross, and visual stimuli. <u>b)</u> and <u>c)</u> represent the close hand <u>(b)</u> and the distant hand condition <u>(c)</u>. In both experiments, the visual stimuli (here displayed as grey circles) were projected one at a time, in one of the ten possible positions (from V-P1 to V-P10), corrected for retinal size <u>(a-c)</u>. Tactile and visual stimuli were presented alone (unisensory) or coupled synchronously with each other (multisensory). Globally, we adopted two conditions of unisensory stimulation (only tactile or visual stimulation) and a multisensory condition (visuo-tactile stimulation). To these, we added catch trials (nor visual nor tactile stimuli presented) to monitor participant’s compliance.

*Tactile stimuli* were brief electrocutaneous stimulations (100 µs, 400 mV) delivered to the right index finger via a constant current stimulator (DS7A, DigiTimer, UK) through a pair of disposable electrodes (1.5*1.9 cm, Neuroline, Ambu, Denmark). Their intensity was determined through an ascending and a descending staircase procedure, incrementing and decrementing, respectively, the intensity of the stimulation to find the minimum intensity at which the participant could detect 100% of the touches over 10 consecutive stimulations. Intensity was further increased by 10% before the first and third experimental block.

#### Design & Procedure

Participants performed a speeded tactile detection task. Tactile stimulation of their right index finger could be delivered alone or synchronous to a visual one, in one of the ten positions (see Figure 1). Participants rested with their head on the chinrest and eyes on the fixation cross. Their right hand was placed on the table 16 cm rightward from the body’s sagittal axis, with the tip of the middle finger in correspondence to V-P2 (hereafter close hand) or V-P6 (hereafter distant hand), in different blocks counterbalanced across participants (116 randomized trials per block): two blocks with the close hand and two with the distant hand. Considering the distance between the positions of visual stimulation, the hand in the distant position covers positions V-4 (wrist), V-5 and V-P6 (tip of the middle finger), as well as the hand in the close position is flanked by the positions V-P1 and V-P2 (see Figure 1). Each hand condition included 16 visuo-tactile (VT) stimulations per position and 16 unimodal tactile trials (T trials). To ensure compliance with task instructions, there were also 4 unimodal visual trials per position (V trials) and 16 trials with no stimulation (N trials). Participants had to respond to the tactile stimulus as fast as possible pressing a pedal with their right foot. The total duration of the experiment was about 45 minutes.

#### Analyses

Both the experiments adopted a within-subject design. When necessary, Green-house-Geysser sphericity correction was applied. The first analyses focused on the accuracy of the performance. Four participants performed poorly (>2 SD from mean) and were excluded from further analyses.

To have a direct index of the proportion of multisensory facilitation over the unimodal tactile condition, we calculated the Multisensory Gain (MG):

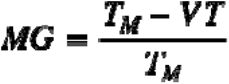

T_M_ was the average RT for unimodal tactile stimuli, and VT was the raw RT for a multisensory visuo-tactile stimulus. Larger MG values correspond to greater facilitation (namely, larger benefits for VT compared to T conditions). This measure is more rigorous than an absolute delta, as it allows correction of the RTs considering the subject-specific speed for each visual position and for each position of the hand (analyses on the delta RT are also reported in Appendix – Experiment I). Computing MG values per hand and stimulus position, we obtained two vectors of 10 MG values (from V-P1 to V-P10) for each participant: one for the close hand and one for the distant hand. We applied a multivariate SVM approach (Vapnick, 1995) to test whether a data-driven classifier could reliably predict the position of the hand from the spatial pattern of MG. The SVM was trained on (N – 1) participants (leave-one-out strategy) and tested on the two vectors excluded from training, using a linear kernel. Overall accuracy was calculated as the sum of the correct predictions for both hand positions divided by the total number of predictions.

To map multisensory facilitation more locally, we compared Bonferroni-corrected MG values for each position against zero and performed a *Hand* (close vs. distant)\**Position* (V-P1 to V-P10) within-subject ANOVA.

To compare the shape of these multisensory facilitation maps, we first tested which function better fit the spatial pattern and, second, we cross-correlated them to test their shapes for isomorphism. MG values were fitted to sigmoidal and normal curves, limited to two parameters. Table 1 reports formulas for curve fitting (Curve Fitting toolbox) with MATLAB (version R2016b, MathWorks, USA). Similar to previous work (Canzoneri et al., 2012; Serino et al., 2015), we considered a good sigmoidal fit when data fitted a descending slope, indicating a facilitation close to the body that fades away with increasing distances.

**Table 1:** Formulas adopted to fit the curves for the multisensory gain values in Experiment I. X represents one of the 10 experimental positions (from V-P1 to V-P10). We used the same formulas to fit the sigmoidal and normal curves to reachability judgements in Experiment II.

| Sigmoidal | Normal |
| --- | --- |
| $\frac{100}{1 + e^{-a(X-h)}}$ | $100 * e^{-\left(\frac{X-a}{b}\right)^2}$ |

Then, we performed a cross-correlation analysis on MG values to evaluate the isomorphism of the facilitation curve for both hand positions. Our prediction was that shifting the close hand pattern of facilitation distally (i.e., towards the distant hand position), should bring to higher correlations due to the overlap of the curves. We correlated the pattern of averaged MG values for all reachable stimuli (V-P1 to V-P6) in the close hand condition, with that of six averaged MG values observed in the hand distant condition. The correlation was then tested for four incremental position shifts (distally, 1 per position), up to the last shift, where we correlated the V-P1 to V-P6 pattern for the close hand with the V-P5 to V-P10 pattern of the distant hand.

### Results

We tested the effect of VT stimulation over 10 uniformly spaced positions, to obtain a fine-grained map of patterns of multisensory facilitation (validated in a pilot study). Participants performed accurately (90% hits, <2% false alarms). First, the multivariate classifier was able to predict the two positions of the hand with an accuracy of 0.72 (33/46 correct classifications), with no bias for one hand position over the other (17/23 and 16/23 for the close and distant hand respectively). This accuracy was significantly higher than chance (one-tailed binomial test p=0.002). Hence, different patterns of multisensory facilitation were associated with different hand positions within the ARS.

A *V-Position*\**Hand* repeated measures ANOVA (Figure 2A) revealed a significant main effect of *V-Position* (F_(5.85,128.71)_=3.52, p=0.003, η^2^_p_=0.14), further modulated by hand position, as indicated by the significant interaction (F_(6.45,141.85)_=3.47, p=0.002, η^2^_p_=0.14). Tukey-corrected multiple t-test comparisons revealed faster responses in V-P2 than in V-P4 and in all the positions from V-P6 to V-P10 when the hand was close (all p_s_<0.05 except V-P2 vs. V-P8, p=0.054); responses were faster in V-P4 than in V-P1, V-P2, V-P3, V-P8, V-P9 and V-P10 when the hand was distant (all p_s_<0.05). Critically, the MG was larger in V-P2 when the hand was close than when it was distant (p=0.041). This pattern was reversed in V-P4, where the MG was larger when the hand was distant than when it was close (p=0.022). No other differences were significant.

**Figure 2:**
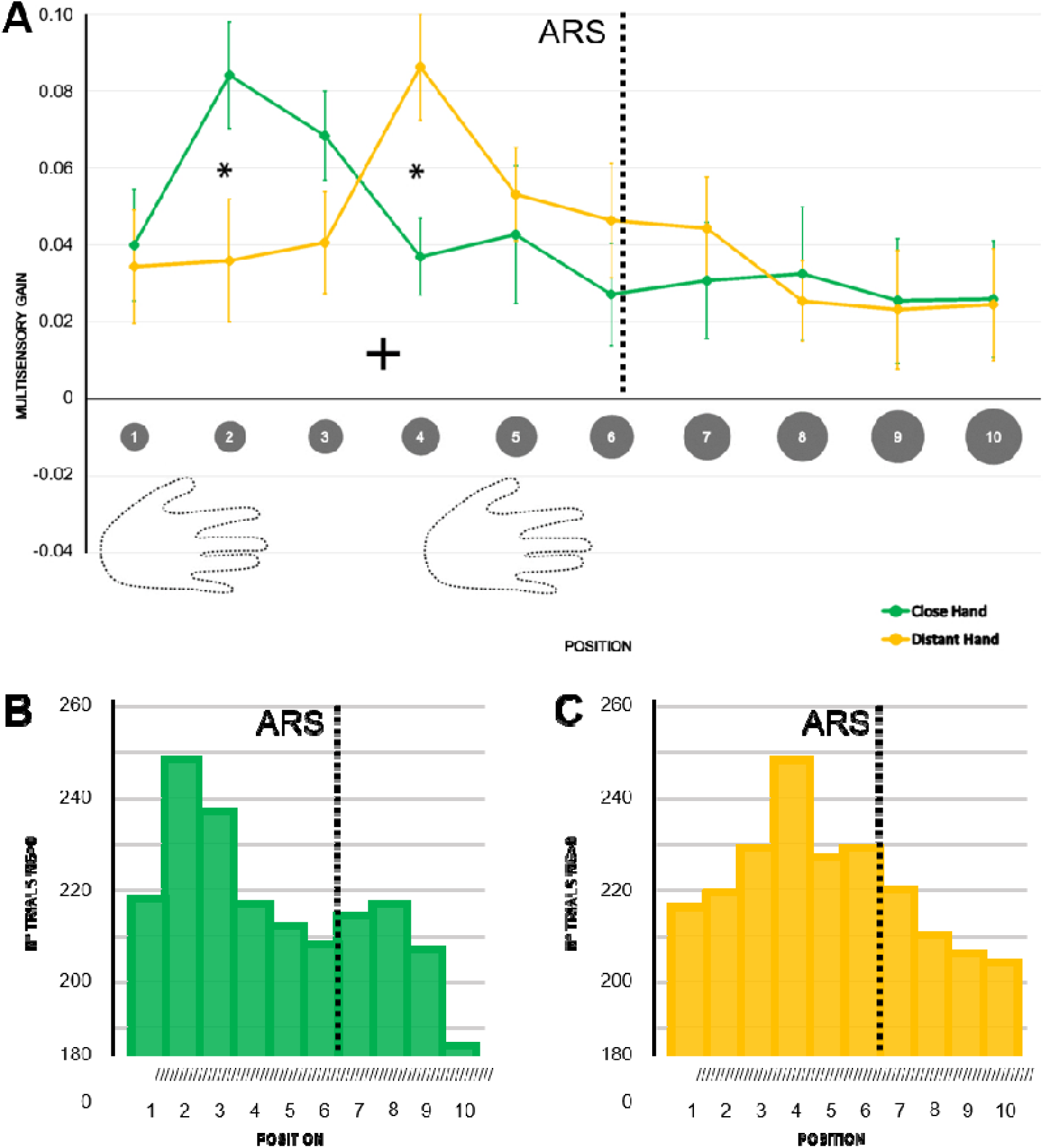
Different patterns of hand-centred multisensory facilitation within ARS. a) Multisensory gain (MG) values along the 10 visual positions, ranging from near to far space, for the distant (yellow) and the close (green) hand conditions. Higher values of MG represent stronger facilitation in terms of RT in the multisensory condition than in the unisensory tactile baseline (by definition, MG = 0). Error bars represent the standard error of the mean. Asterisks represent a significant difference (p < 0.05, corrected). b) and c) Number of trials reporting MG values greater than zero (unisensory tactile baseline) along the 10 visual positions, ranging from near to far space, for the close (b) and the distant (c) hand conditions.

To identify where multisensory facilitation was significant at the single position level, we ran a series of Bonferroni-corrected t-tests on the MG values vs. 0 (i.e., no facilitation). When the hand was close, the MG significantly differed from 0 in V-P2 and V-P3 (all ps<0.05). In contrast, when the hand was distant, the MG was larger in V-P4, V-P5 (all ps<0.05) and marginally in V-P6 (p =0.055). Figure 2 shows the number of trials reporting MG values greater than 0 with the hand close (2B) and distant (2C). The density peak shifted coherently with the position of the hand within ARS. Similar results were obtained by analyzing the delta RT for both the ANOVA and the t-tests (see Appendix–Experiment I). Furthermore, the results of Experiment S1 (see Appendices-S1) show that this multisensory facilitation does not depend on sheer attentional factors.

These findings highlight the hand-centred nature of the multisensory facilitation, occurring in different locations, depending on hand position. From this, one would expect 1) the facilitation to be maximal in correspondence with hand location and to decay with distance from it and 2) the bell-shaped pattern of facilitation to follow the hand when it changes position. To test the first prediction, we modelled our data to a Gaussian curve. To test the alternative hypothesis, namely, that facilitation spreads all over the ARS to decay when approaching the reachable limit, we compared the Gaussian to a sigmoid function fitting (Canzoneri et al., 2012; Serino et al., 2015). The sigmoidal curve could fit the data for a limited number of participants (distant hand: 5/23 subjects, 21.7%; close hand: 9/23 subjects, 39.1%). Instead, fitting the Gaussian curve to the same data accommodated convergence problems for a higher number of participants (distant hand: 14/23 subjects, 59.9%; close hand: 15/23 subjects, 65.2%).

The second prediction, that the bell-shaped facilitation should shift following the hand, was confirmed by the estimation of the position of the peak of the Gaussian curve in each hand position: with the hand close, the peak fell between V-P2 and V-P3 (2.34±1.51); with the hand distant, it fell between V-P4 and V-P5 (4.15±1.28). We then performed a cross-correlation analysis testing whether the curves reported for the two hand positions overlapped when considered in absolute terms. We reasoned that shifting the position of the hand–within the ARS–should bring to an isomorphic facilitation around the new hand position. This would imply the maximum correlation between MG values to emerge when the close-hand curve shifts distally, towards the distant-hand position curve. We considered the first six values of MG with the close hand (from V-P1 to V-P6, i.e., the reachable positions) and correlated this distribution with six values of the MG for the distant hand (Figure 3). We found the maximum correlation (r=0.94 p=0.005) when shifting the close hand distally by two positions. No other correlations were significant (all p_s_>0.20).

**Figure 3:**
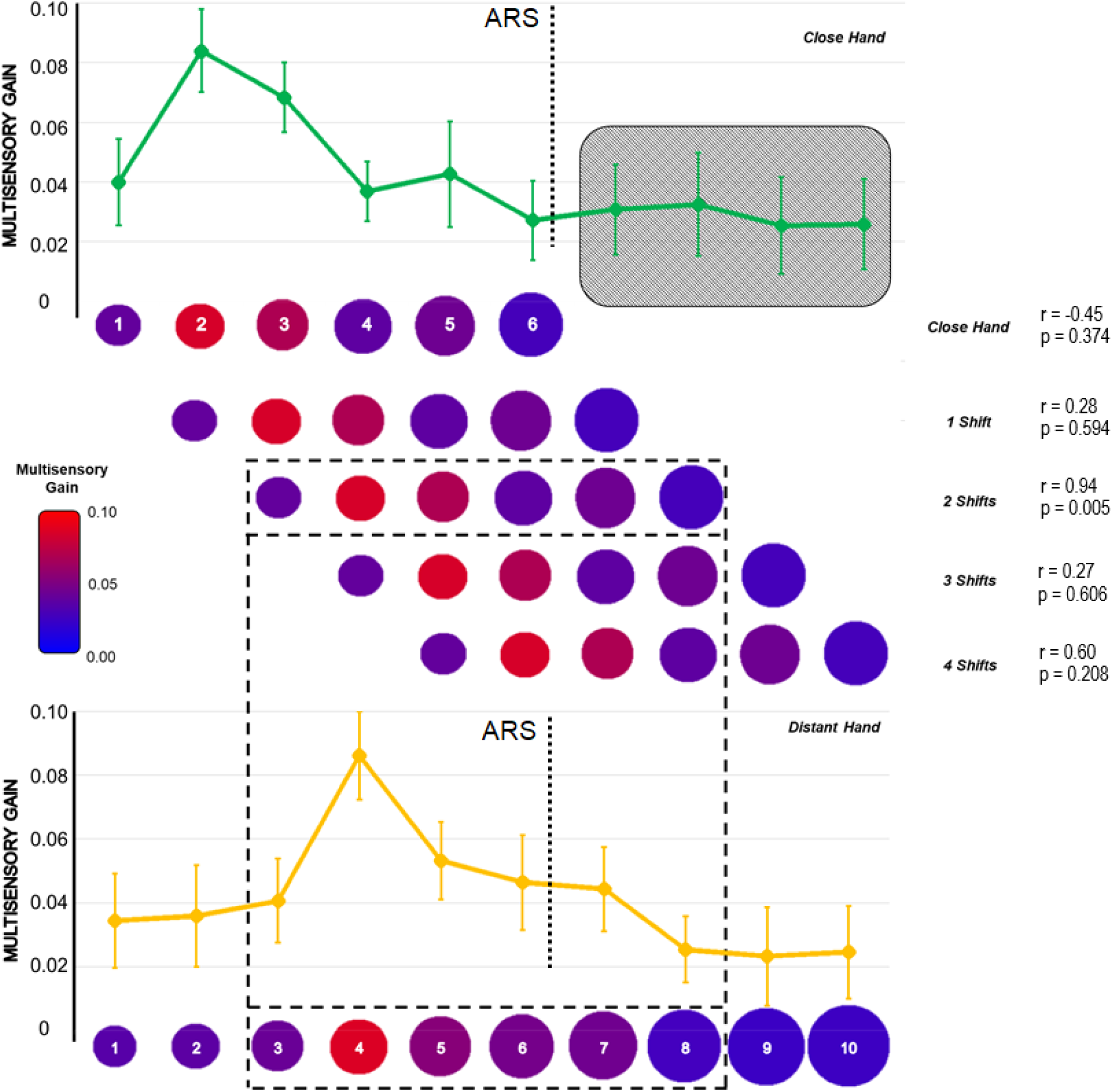
The spatial pattern of MG shifts and follows the hand within reaching space. Cross-correlation analysis of distally shifting the pattern of MG values for all reachable positions with the hand close. Red colors represent higher MG values. Values of Pearson’s r and p values are reported for all the correlations performed. The black grid highlights the only significant correlation (p < 0.05).

### Discussion

The results of Experiment I clearly indicate that PPS and ARS are not superimposable. Yet we cannot exclude that a reachability judgement task might still capture some of the PPS features. To investigate this possibility, we performed three experiments adopting this task and the same settings of Experiment I. The results of Experiments S2 and S3 replicated well-established findings about the ARS, including the overestimation of its limit (Bourgeois & Coello, 2012; Carello et al., 1989). However, they failed to show any similarity with PPS, either in terms of absolute extent (ARS is larger than PPS) or position-dependent modulation (PPS is hand-centred, whereas ARS is not, see Appendices-S2 and S3). To allow a full comparison, in Experiment II we made the reachability judgment task as similar as possible to the tactile detection task, using the same hand positions and multisensory stimulations.

## Experiment II

### Methods

#### Participants

Twenty-five (16 females; mean age=24.44, range=18-41; mean arm length=78.46±7.26 cm) participants matching the same criteria as Experiment I participated in Experiment II.

#### Stimuli and Apparatus

*Visual* and *tactile* stimuli were identical to those used in Experiment I.

#### Design & Procedure

We took advantage of an ARS multisensory task by asking participants to perform reachability judgements while tactile stimuli were concurrently presented with the visual stimulus. Experiment II was meant to assess whether the multisensory stimulation (in addition to having the hand visible and in the same positions as Experiment I) could either induce hand-centred facilitation in the reachability task performance and/or change the extent of the reachability limit. We employed the same settings of Experiment I and applied the same tactile stimulation to the right index finger, placed in either the close or the distant position. However, in this case the tactile stimulus was task-irrelevant. Overall, 160 randomized V and 160 randomized VT trials were presented for each hand position, administered in two blocks in a randomized order. The order of hand positions was counterbalanced across participants.

#### Analyses

Similar to Experiment I, we tested the classifier on the MG patterns and performed the same procedures already described on delta RTs and MG. The percentage of “reachable” responses per position was calculated and then fitted to sigmoidal and normal curves, as in Experiment I. We fitted the curves separating hand positions and type of stimulation (unimodal visual versus multisensory visuo-tactile). *Hand* (close vs. distant)\**Stimulation* (visual vs. visuotactile)\**Model* (Gaussian vs. Sigmoid) ANOVA on RMSE values assessed which model best fitted the data, both at the individual and group level. Either way, the best-fitting model for these data was the sigmoidal curve. Thus, we investigated the PSE and slope values by subjecting them to two separate repeated measure ANOVAs with *Hand* (close vs. distant) and *Stimulation* (visual vs. visuo-tactile) as within-subject factors.

### Results

Participants were accurate (>90% hits, <2% false alarms). We computed for each subject two vectors of MG values, as in Experiment I, and we could leverage a similar data-driven classifier to discriminate the close from the distant hand. Prediction accuracy was lower than in Experiment I (0.36, 18/50 correct classifications) and not significantly higher than chance level (one-tailed binomial test p=0.98), indicating that the classifier failed to distinguish between hand positions within ARS.

Moreover, the V-*Position*\**Hand* within-subject ANOVA on the MG did not reveal any significant effect (Hand: F_(1,24)_=0.83, p=0.37; V-Position: F_(6.31,151.56)_=1.20, p=0.31; Hand*V-Position: F_(5.35,128.5)_=1.82, p=0.11). However, the significant intercept (F_(1,24)_=9.80, p=0.005) confirmed the general facilitation produced by multisensory stimulation, with respect to the unisensory one. Multiple Bonferroni-corrected comparisons revealed that none of the positions presented an MG significantly different from 0 (all p_s_>0.05) when the hand was close. V-P5 and V-P6 differed from 0 (all p_s_<0.05) when the hand was distant. Similar results were obtained by analyzing the delta RT, both with ANOVA and t-test (see Appendix–Experiment II).

Reachability judgements were then fitted to sigmoidal and Gaussian curves. Within-subject ANOVA on the RMSE of these models was performed with a *Model* (sigmoidal vs. Gaussian)\**Stimulation* (visual vs. visuo-tactile)\**Hand* (close vs. distant) design. The sigmoidal curve reported the best fit, irrespective of stimulation type and hand position (Model: (F_(1,24)_=220.11, p<0.001, η^2^_p_=0.90)). For each variable, we estimated the coefficients of the sigmoid, obtaining the PSE and the curve slope. Through a *Hand* (close vs. distant)\**Stimulation* (visual vs. visuo-tactile) within-subject ANOVA on PSE values, we observed a main effect of stimulation type (F_(1,24)_=4.38, p=0.05, η^2^_p_=0.15): the mean PSE was closer to the body in the unimodal visual (mean±SE = 6.67±0.17) than in the multisensory visuo-tactile condition (6.76±0.18). The main effect of Hand (F_(1,24)_=0.07, p=0.79) and its interaction with Stimulation (F_(1,24)_=3.49, p=0.07) were not significant. Last, we performed a *Hand* (close vs. distant)\**Stimulation* (visual vs. visuo-tactile) within-subject ANOVA on slope values. Neither main effects (Hand: F_(1,24)_=1.75, p=0.20; Stimulation: F_(1,24)_=0.35, p=0.56) nor the interaction (Hand*Stimulation: F_(1,24)_=0.27, p=0.61) were significant (Figure 4).

**Figure 4:**
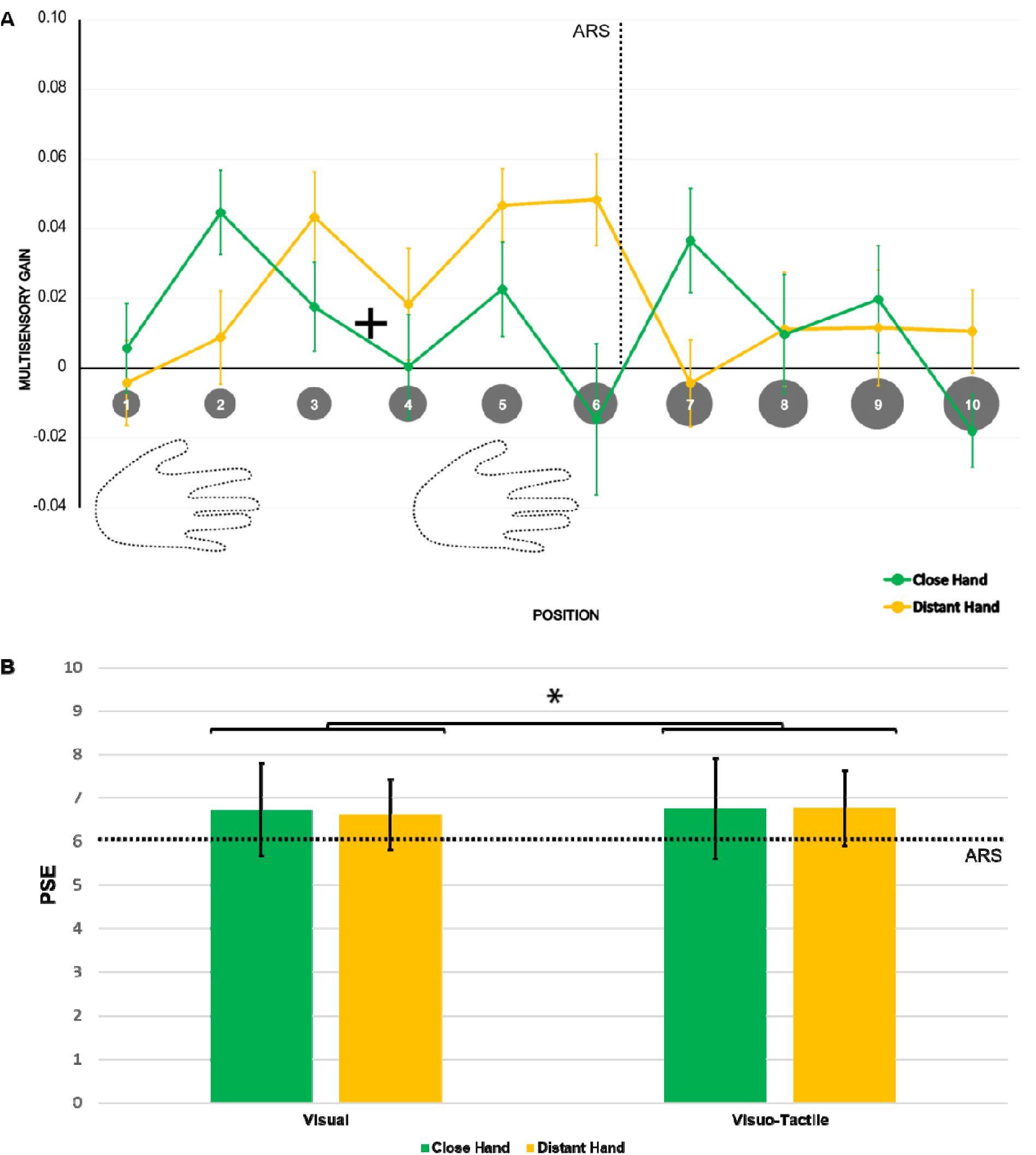
No hand-centred MG spatial patterns in a reachability judgement task. a) Multisensory gain (MG) values along the 10 positions, ranging from near to far space, for the close (green) and distant (yellow) hand conditions. Higher values of MG represent a stronger facilitation in terms of RT with respect to the unimodal visual baseline (by definition, MG = 0). Error bars represent the standard error of the mean. No significant differences between hand postures emerged. b) PSE values calculated for both unimodal visual and multisensory visuo-tactile conditions for both hands. Error bars represent the standard error of the mean. Asterisks indicate a significant difference between unisensory and multisensory conditions (p < 0.05).

## Discussion

We contrasted two theoretical views about PPS and ARS: one proposing they are different, the other opposing they are the same. Our findings clearly point against the latter, whether contrasted in terms of their spatial extent, by using their respective gold-standard paradigms and measures, or in terms of pattern of multisensory facilitation.

Due to obvious differences between paradigms, we did not compare the multisensory facilitation directly. We rather reasoned that, would PPS and ARS be the same spatial representation, using their typical paradigms applied to the same body-part we should obtain similar results. Our visuo-tactile version of the reachability judgement task confirms previous findings on the extent and overestimation of ARS (Bootsma et al., 1992; Bourgeois & Coello, 2012; Carello et al., 1989; Coello & Iwanow, 2006), but its comparison with the PPS multisensory task resulted in two main advances arguing against the PPS-ARS identity.

First, we observed that multisensory facilitation depends on hand position, peaking in correspondence with its location and decaying with distance from it. Notably, this near-hand facilitation effect is independent of attention orienting (see S1 results). Thus, PPS is smaller than the ARS, either objectively (from V-P1 to V-P6) or subjectively (PSE) measured. Were they superimposable, we should have observed faster RTs for all the reachable positions of visual stimulation. Both the classifier and the location-specific differences indicate instead that different spatial patterns of multisensory facilitation emerge for the close and distant hand positions, despite being both within the ARS limits. Interestingly, we add that overestimation is not modulated by hand vision (see Experiments S2 and S3), and is independent of the position of the hand (Experiments S3 and II).

Second, our findings indicate that ARS is not hand-centred, whereas PPS is. In Experiment II, adapting the reachability judgement task to a multisensory setting, the only significant effect was a general multisensory facilitation, spread over the 10 positions tested: there was no modulation as a function of stimulus reachability or hand proximity which, on the contrary, define PPS (Experiment I). Therefore, ARS is not encoded in a hand-centred reference frame. Indeed, hand position was robustly classified from the distribution of MG in Experiment I (PPS), but not in Experiment II (ARS). Thus, the proximity of visual stimuli to the hand –not their reachability– predicts the increase in multisensory facilitation. Cross-correlation and univariate analyses further demonstrated that visual boosting of touch is hand-centred, following changes in hand position. In sum, here we show that 1) PPS does not cover the entire ARS, 2) ARS is not hand-centred, and 3) ARS is not susceptible to multisensory stimulation. Taken together, these results converge in showing that PPS and ARS are not superimposable. Previous neuroimaging (Brozzoli et al., 2011, 2012) and behavioral studies (di Pellegrino et al., 1997; Farnè et al., 2005; Serino et al., 2015) reported body-part centered multisensory facilitation within PPS. Here we disclose that the facilitation is isomorphically “anchored” to the hand: present in close positions when the hand is close, it shifts to farther positions when the hand is distant, without changing its ‘shape’. Notably, the facilitation pattern fits well a Gaussian curve, similar to what observed in non-human primate studies (Graziano et al., 1997) and in line with the idea of PPS as a « field », gradually decaying around the hand (Bufacchi & Iannetti, 2018).

The amount of multisensory facilitation observed in Experiment I for the position closest to the trunk (V-P1, thus clearly within ARS) is also remarkable. First, it is lower than that observed in correspondence of the close-hand PPS peak (between V-P2 and V-P3) and, second, it is comparable to that obtained for all the out-of-reach positions (V-P7 to V-P10), irrespectively of hand distance.

These findings are consistent with what one would predict from neurophysiological data. Studies on non-human primates requiring reaching movements performed with the upper limb found activations involving M1, PMv and PMd, parietal areas V6A and 5 and the parietal reach region (Buneo et al., 2016; Caminiti et al., 1990; Georgopoulos et al., 1982; Kalaska et al., 1983; Mushiake et al., 1997; Pesaran et al., 2006). In humans, ARS tasks require judging stimulus reachability (Carello et al., 1989; Coello et al., 2008; Coello & Iwanow, 2006; Rochat & Wraga, 1997) or performing reaching movements (Battaglia-Mayer et al., 2000; Caminiti et al., 1990, 1991; Gallivan et al., 2009). Brain activations underlying these tasks encompass M1, PMd, supplementary motor area, posterior parietal cortex and V6A as well as the anterior and medial IPS (Lara et al., 2018; Monaco et al., 2011; Pitzalis et al., 2013; see Filimon, 2010 for review). Therefore, despite some overlap in their respective fronto-parietal circuitry, PPS and ARS networks do involve specific and distinct neuroanatomical regions, in keeping with the behavioral differences reported here.

At odds with previous studies employing looming stimuli (Canzoneri et al., 2012; Finisguerra et al., 2015; Noel et al., 2015; Serino et al., 2015, but see Noel et al., 2020), we used ‘static’ stimuli flashed with tactile ones to avoid inflating the estimates of multisensory facilitation. Looming stimuli with predictable arrival times induce foreperiod effects that, though not solely responsible for the boosting of touch, may lead to overestimations of the magnitude of the facilitation (Hobeika et al., 2020; Kandula et al., 2017). Most noteworthy, the findings of the attentional control experiment provide the first behavioral evidence that multisensory near-hand effects may be appropriately interpreted within the theoretical framework of peripersonal space coding. This study therefore offers a bias free (Holmes et al., 2020) protocol for fine-grained mapping of PPS.

In conclusion, this study provides an empirical and theoretical distinction between PPS and ARS. Discrepancies concern both their spatial extent and their behavioral features, and warn against the fallacy of conflating them. A precise assessment of PPS is crucial because several researchers exploit its body-part-centred nature as an empirical entrance to the study of the bodily self (Blanke et al., 2015; Noel et al., 2015; Makin et al., 2008). Moreover, our results have direct implications for the study of interpersonal space, defined as the space that people maintain with others during social interactions. Several studies drew conclusions about interpersonal space using reachability tasks (Cartaud et al., 2018; Iachini et al., 2014; Bogdanova et al., under review). The present findings make clear that using these tasks does not warrant any conclusion extending to PPS, or informing about its relationship with the interpersonal space. Instead, they highlight the need to investigate the potential interactions between PPS and ARS, as to better tune rehabilitative protocols or brain machine interface algorithms for the sensorimotor control of prosthetic arms, for which multisensory integration appears crucial (Makin et al., 2007; Suminski et al., 2009, 2010).

## Acknowledgements

This work was supported by the Labex/Idex (ANR-11-LABX-0042) and by grants from the James S. McDonnell Foundation, the ANR-16-CE28-0015-01 and ANR-10-IBHU-0003 to A.F. E.B. was supported by the European Union’s Horizon 2020 research and innovation programme (Marie Curie Actions) under grant agreement MSCA-IF-2016-746154; a grant from MIUR (Departments of Excellence DM 11/05/2017 n. 262) to the Department of General Psychology, University of Padova. C.B. was supported by a grant from the Swedish Research Council (2015-01717) and ANR-JC (ANR-16-CE28-0008-01). We thank S Alouche, JL Borach, S Chinel, A Fargeot, S Terrones for administrative and informatics support and F Volland for customizing the setup.

## Open Practices Statements

The data and materials for all experiments are available online at https://osf.io/8dnga/?view_only=b0a4a90d5a8a43bb8ab787b1adf7d5c0

## Appendix

### Experiment I

We calculated the *delta RT,* obtained as the difference between the mean tactile RTs (baseline) for the specific condition (close or distant hand) and visuo-tactile RTs. Averaging the results per condition, we obtained a delta value for each visual position, from V-P1 to V-P10, for each hand posture. Larger values of *delta RT* indicate faster responses. We thus compared these deltas to zero (i.e., the absence of multisensory facilitation) correcting the multiple tests for Bonferroni correction. These comparisons highlighted a significant facilitation from V-P4 to V-P6 (all p_s_ < 0.05) when the hand was placed in a distant position, with a marginally significant difference in V-P7 (p = 0.06), whereas moving the hand to the close position resulted in a significant facilitation only in V-P2 and V-P3 (all ps < 0.05). Subsequently, we performed a *Hand* (close vs. distant) * *Position* (V-P1 to V-P10) within-subject ANOVA on these delta values to compare the multisensory facilitation obtained at each hand position and to obtain a measure of the extent of this facilitation (namely, the extent of the hand-centred PPS, see Figure S1). Moreover, between-hand position comparisons allowed us to observe whether the position and the extent of this PPS-related facilitation moved in a hand-centred fashion. The main effect of Hand was not significant (F_(1,22)_ = 0.01, p = 0.94), whereas the main effect of Position was significant (F_(5.86,129.02)_ = 3.50, p = 0.003, η^2^_p_ = 0.14). However, this effect was modulated by hand location (significant Hand*Position interaction, F_(6.60,145.23)_ = 3.39, p = 0.003, η^2^_p_ = 0.13). Tukey-corrected multiple comparisons within position revealed significantly faster responses in V-P2 versus V-P4, V-P6, V-P7, V-P9 and V-P10 for the close hand (all p_s_ < 0.05), with a marginally significant difference between V-P2 and V-P8 (p = 0.060); for the distant hand, V-P4 resulted in significantly faster responses than V-P1, V-P2, V-P3, V-P8, V-P9 and V-P10 (all p_s_ < 0.05). Comparing the hand positions, we observed greater facilitation in V-P2 with the close hand than with the distant hand (p = 0.053) and the opposite pattern was observed in V-P4 (p = 0.037). No other differences were significant. Due to the limited number of participants who reported a good fit with the sigmoidal curve on MG values, it was not possible to directly compare the RMSE values obtained with those concerning the fitting of a Gaussian curve. We thus performed a chi-square test with Yates’ continuity correction (Yates, 1934) on the percentages of best fit cases for the distant hand (21.7% for the sigmoidal curve, 59.9% for the Gaussian one), revealing that the Gaussian model fit for a larger proportion of subjects than the sigmoid model (χ^2^_(1)_ = 5.739, unilateral p = 0.008). The same was true with the hand in the close position (39.1% for the sigmoidal curve, 65.2% for the Gaussian one), though the difference was marginally significant χ = 2.178, unilateral p = 0.069). This may be due to the number of visual positions available in the space preceding the peak of facilitation with the hand close. Indeed, with the hand placed in the distant location, several visual positions before and after the peak of facilitation were present, providing optimal conditions for testing both Gaussian and sigmoidal models. By contrast, with the hand placed close, only one stimulus position was available before the peak, providing equally optimal conditions for the sigmoidal model as the hand distant condition but non-optimal conditions for the Gaussian model. It is possible that this configuration of the setting influenced the fit by biasing sigmoidal over Gaussian fitting in the condition with the hand close. Despite this, however, the Gaussian curve still resulted as the best fitting for most participants.

**Figure S1.**
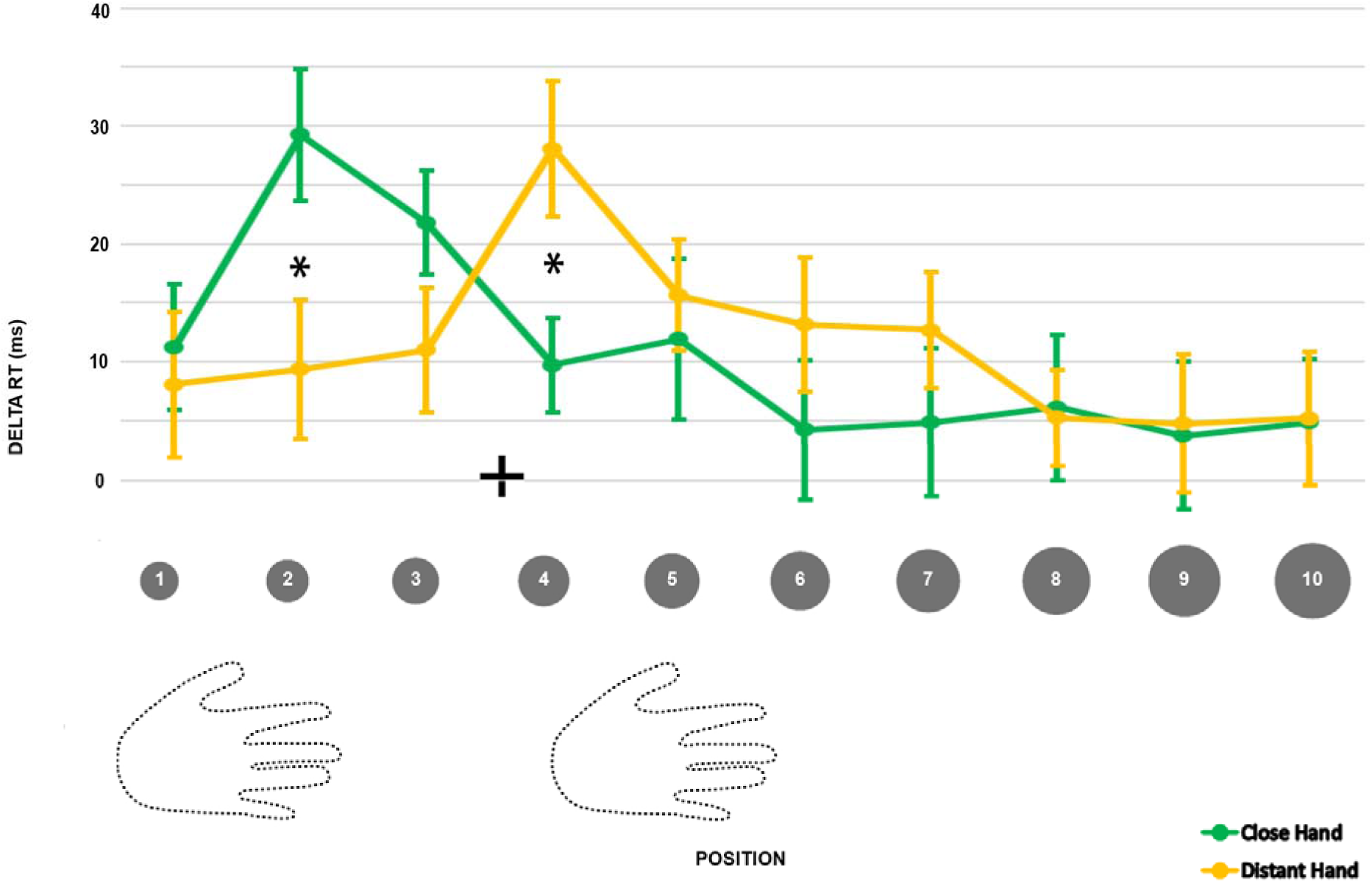
Delta RT values along the 10 visual positions, ranging from near to far space, for the distant (yellow) and the close (green) hand conditions. Higher values of delta RT represent a stronger facilitation in terms of RT for the multisensory visuo-tactile condition than for the unisensory tactile baseline (by definition, delta RT = 0). Error bars represent the standard error of the mean. Asterisks represent a significant difference (p < 0.05, corrected).

### Experiment II

We calculated the delta RT as done in Experiment I. Thus, we compared the averaged delta for each position to 0 (namely, the absence of facilitation). Bonferroni-corrected multiple comparisons reported only a marginally significant facilitation in V-P2 in the close hand condition (p = 0.058), whereas significant facilitations emerged in V-P3, V-P5 and V-P6 with the distant hand (all p_s_ < 0.05). This was the first main difference from Experiment I, in which a facilitation emerged with both hand positions.

We subjected these values to a *Hand* (close vs. distant) * *Position* (V-P1 to V-P10) within-subject ANOVA to compare the facilitation obtained in the same visual positions with different hand locations. Neither the main effect of Hand (F_(1,24)_ = 0.56, p = 0.46) nor the effect of Position (F_(6.56,157.38)_ = 1.17, p = 0.33) were significant. In the same way, the Hand*Position interaction was non-significant (F_(5.42,130.19)_ = 2.01, p = 0.08). Therefore, Experiment II only highlighted a general effect of the multisensory stimulation over the unimodal visual stimulation, as reported by the significant intercept (F_(1,24)_ = 14.91, p < 0.001); however, this effect was independent of the position of the visual stimulation and was not hand-centred. These results are reported in Figure S2.

**Figure S2.**
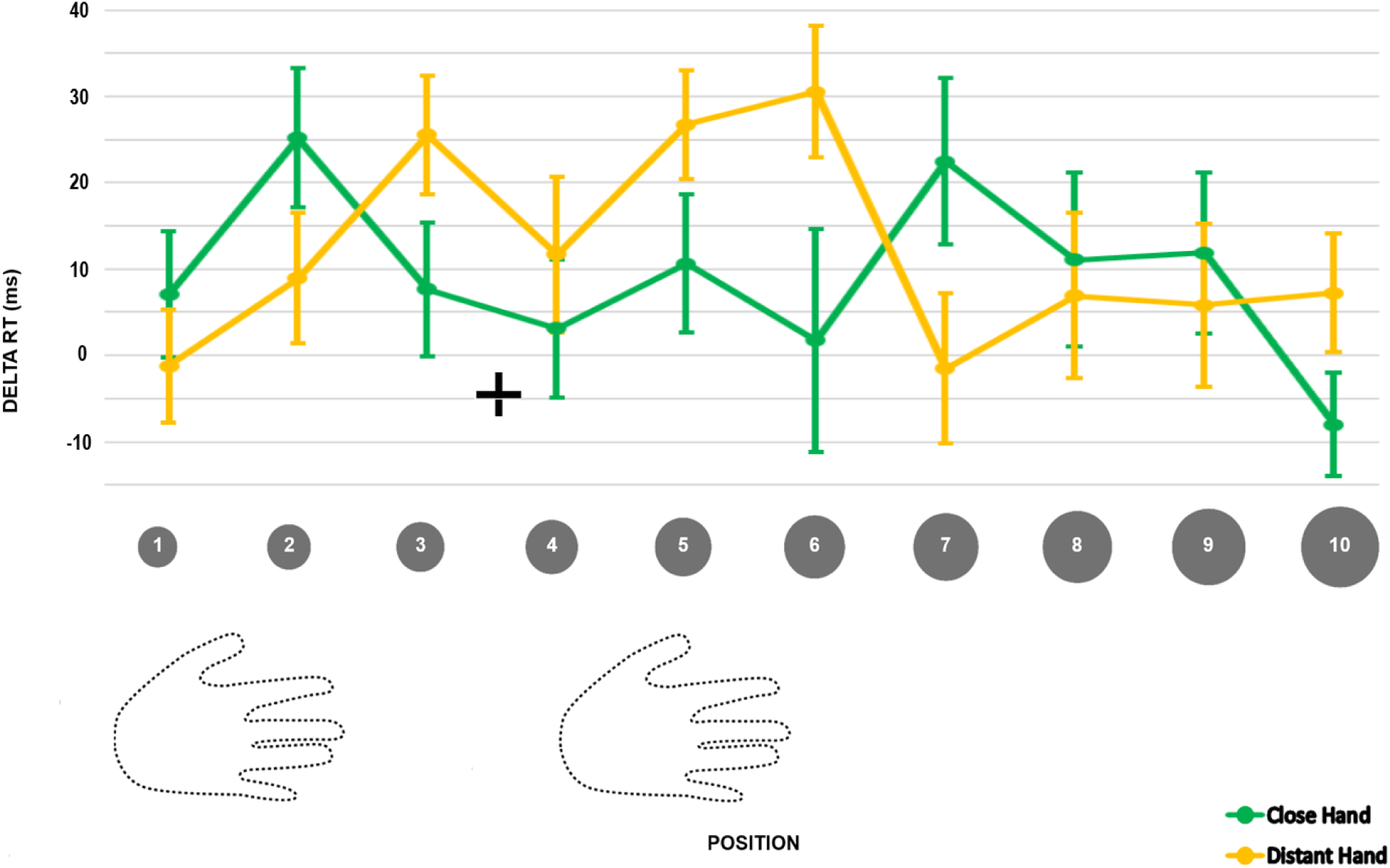
Delta RT values along the 10 visual positions, ranging from near to far space, for the distant (yellow) and the close (green) hand conditions. Higher values of delta RT represent a stronger facilitation in terms of RT for the multisensory visuo-tactile condition than for the unisensory tactile baseline (by definition, delta RT = 0). Error bars represent the standard error of the mean.

Starting from the raw reachability judgments, we calculated for each participant the percentage of “reachable” responses per position of visual stimulation, separating the type of stimulus (visual or visual-tactile) and the hand position (close or distant). We then subjected these percentages to an ANOVA 2 (Hand: close or distant) * 2 (Condition: V or VT) * 10 (Visual position, from V-P1 to V-P10). We observed that the main effect of Position is significant: F_(1.53,36.69)_ = 331.63, p < .001, η^2^_p_ = 0.93. The post-hoc with Bonferroni correction show a clear difference between the judgments relating to the first five positions (not statistically different between them, all p_s_ > 0.05) and those relating to the last five (all p_s_ > 0.05). Since the position V-P6 is the objective limit of participants’ reachable space, these results therefore seem to confirm that the subjective judgment becomes less secure at this limit. Moreover, the interaction Hand*Condition is significant: F_(1,24)_ = 7.13, p = 0.013, η^2^_p_ = 0.23. Bonferroni-corrected post-hocs show a difference between the reachable response rate of the unisensory visual condition (mean ± se = 61.1 ± 1.65) and the visual-tactile condition (62.8 ± 1.59), but only when the hand is in the far position (p = 0.004). Even if the absence of a significant interaction with the visual stimulation position prevents us from verifying it directly, we believe it is legitimate to hypothesize that this difference can be related to that found in terms of PSE in the modeling analyses of these judgments. A PSE located at a greater distance in the visual-tactile condition, in fact, involves a greater percentage of “reachable” responses in this condition in the positions closest to the objective limit of reaching. No other significant effect emerged. The percentage of reachability judgments is reported in Figure S3.

**Figure S3.**
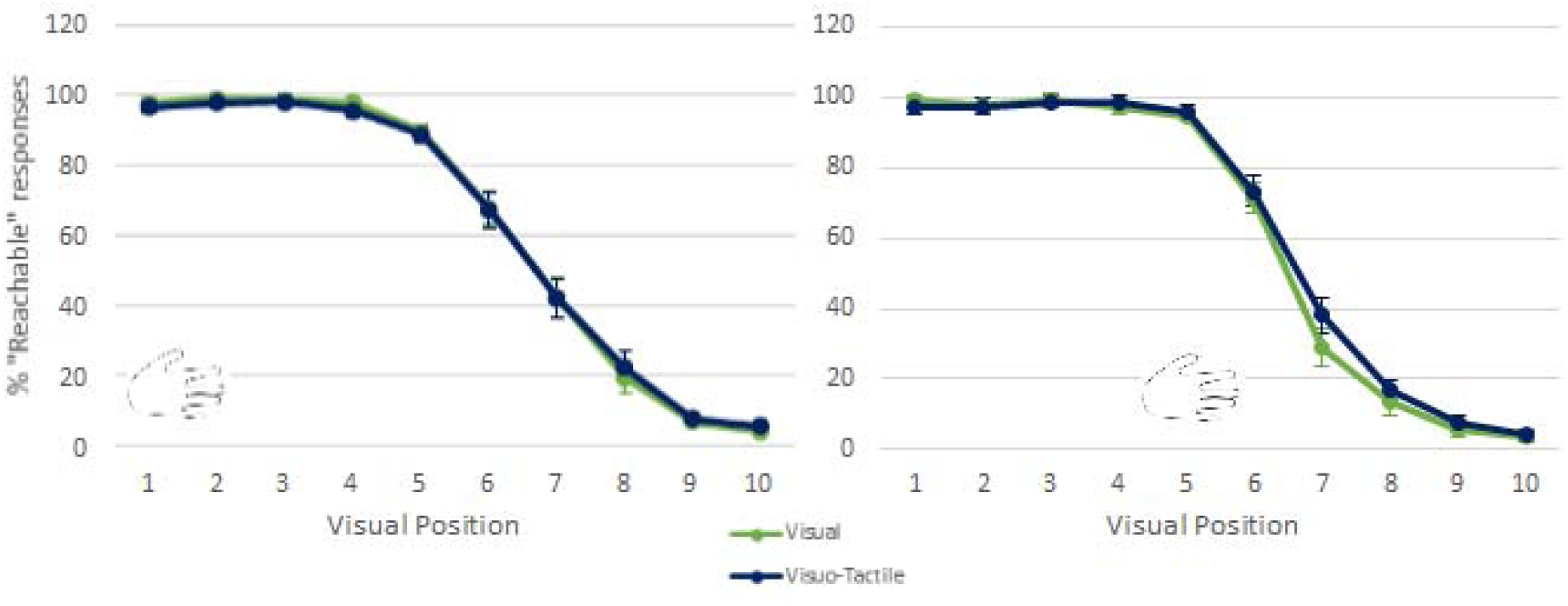
Percentages of reachability judgments for the unisensory visual (green) and multisensory visuo-tactile (blue) conditions along the 10 positions of visual stimulation for the close (left panel) and distant hand (right panel). The 6^th^ visual position represents the objective reachable limit. Error bars represent the standard error of the mean.

### Experiment S1

The hand-centered multisensory facilitation described in Experiment I could contain some contribution of attention: it is possible that a greater attentional focus towards a specific position of visual stimulation could influence performance, accelerating the response. Although this possibility has already been ruled out in the literature (Makin et al., 2009), we ran the Experiment S1 in order to test for the presence of hand-centered facilitation also in case when the attentional focus is shifted away from the hand. As illustrated in Figure S4A-B, we asked our participants to place their right hand on the table, in the same position adopted in Experiment I for the near-hand condition, and to perform a go/no go tactile detection task. Tactile and visual stimuli were identical to those of Experiment I, but in this case we took advantage of only two positions of visual stimulation, thereafter “V-Close” (corresponding to V-P2 of Experiment I) and “V-Distant” (corresponding to V-P5 + 3 cm), equidistant from the fixation cross (15 cm closer and farther respectively). In this way, we were able to exploit the position of visual stimulation that most of all facilitated the performance with the close hand in Experiment I and we opposed a far visual stimulation at the same distance from the position of the gaze and within the reaching space. The diameter of the visual stimulations was corrected for retinal size as a function of their distance, established according to the length of the participant’s right arm, as in Experiment I. We then projected a circular ring (original diameter 3 cm, corrected for retinal size, duration 50 ms) 8 cm leftward to the fixation cross 150 ms before the tactile stimulus. This ring could be completely closed or show a small opening in one of four possible positions (randomized order). We asked our participants to keep their eyes fixed on the fixation cross and to respond as quickly as possible to tactile stimulation, but only if the ring in the left hemispace was closed. Therefore, to ensure good performance, subjects had to shift their attention to the left, while the target and visual stimuli were presented on the right. Being at the same distance from the fixation cross and the ring, neither of the two visual stimuli could be attentively favored over the other.

We established our sample size through an a priori power analysis (G*Power 3.1.9.2), hypothesizing a power of 0.85 and α = 0.05. We thus performed a paired t-test and a Pearson’s correlation between the subject-averaged multisensory gain in V-P2 and V-P5 in the close-hand condition of Experiment I to obtain the effect size (d = 0.71) and the correlation between measures (r = 0.65). Consequently, we needed to recruit at least 20 participants in our Experiment S3 (N = 20, 13 females; mean age = 27.9, range = 22-53; mean arm length = 78.3 cm).

According to Experiment I, each participant was presented with 16 repetitions for each condition (unisensory tactile, close visuo-tactile and distant visuo-tactile) with a “go” signal. To these were added 8 trials per condition with the “no go” signal and 20% of catch trials (visual unisensory stimulation or no stimulation), equally divided between “go” and “no go” signals.

Participants were globally accurate (mean accuracy on “go” trials > 95%, with no difference between the three conditions: F_(1.94, 36.84)_ = 0.76 p = 0.47) and paid attention to the visual stimulus projected on the left (only 6 responses in “no go” trials, 1.25%). To be consistent with Experiment I, only the participants with a mean accuracy greater than 70% (average accuracy of the conditions of Experiment I considered) were considered for subsequent analyses (no participant excluded). RTs greater than 2.33 SD from the participant’s mean in each condition were filtered out as outliers (2.6%). We thus calculated the delta RT as already described in Experiment I and we performed a paired t test between subject-averaged delta values for the Close and the Distant visual conditions. As reported in figure S4C, the difference was significant: t_(19)_ = 2.35, p = 0.030, Cohen’s d = 0.54, indicating faster responses to touches when the irrelevant visual stimulus was presented close to the hand (mean ± se = 22.58 ± 4.02 ms) compared to far from it (4.22 ± 4.96 ms). Coherently with the formula illustrated in Experiment I, we calculated the multisensory gain per participant and per condition, comparing the facilitation obtained in the two visual conditions through a paired t test. As for the delta RT, the difference was significant: t_(19)_ = 2.34, p = 0.031, Cohen’s d = 0.54. In line with our hypothesis, in fact, the facilitation of the performance linked to multisensory stimulation is greater when the visual stimulus is presented near (mean ± se = 0.035 ± 0.008) rather than far from the hand (0.005 ± 0.008).

These results therefore demonstrate that the effects observed in Experiment I cannot be due only to attentional factors, as they persist even after a shift of attention to a point equidistant from the stimulations used and far from the hand.

**Figure S4.**
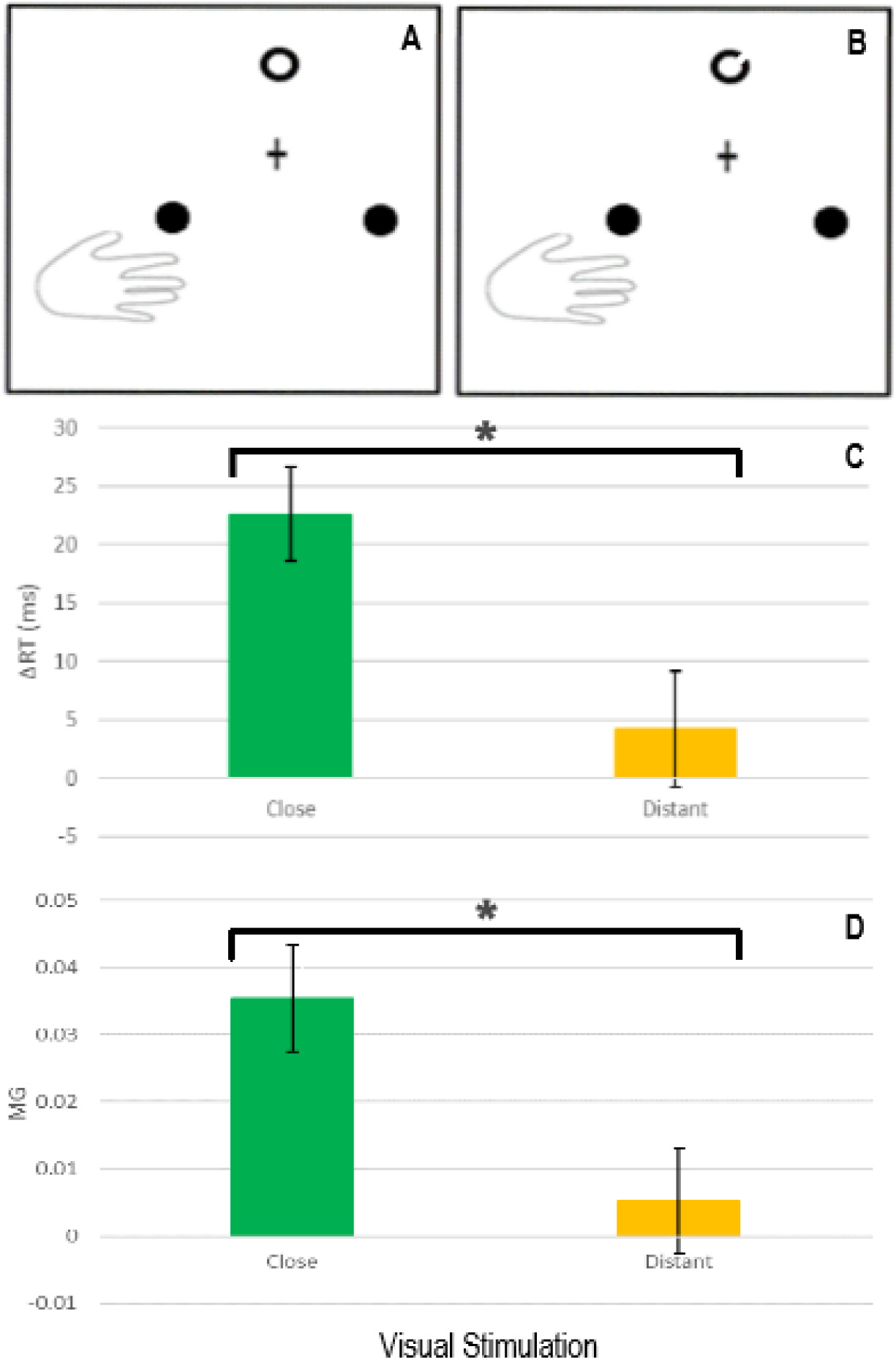
Ruling out attentional factors. A-B) Positions of right hand, fixation cross and visual stimuli Visual stimuli (here displayed as black circles) were projected one at a time, in one of the two possible positions (Close or Distant), corrected for retinal size. Tactile and visual stimuli were presented alone (unisensory) or coupled synchronously with each other (multisensory). Leftward to the fixation cross, a close (A) or open (B) ring represented the “go” and “no go” conditions, respectively. *C)* Delta RT values for the Close (green) and Distant (yellow) visual stimulations. Higher values of delta RT represent a stronger facilitation in terms of RT for the multisensory visuo-tactile condition than for the unisensory tactile baseline (by definition, delta RT = 0). Error bars represent the standard error of the mean. Asterisks represent a significant difference (p<0.05). *D)* Multisensory gain (MG) values for the Close (green) and Distant (yellow) visual stimulations. Higher values of MG represent a stronger facilitation in terms of RT for the multisensory visuo-tactile condition than for the unisensory tactile baseline (by definition, MG = 0). Error bars represent the standard error of the mean. Asterisks represent a significant difference (p<0.05).

### Experiment S2

Experiment S2 aimed to extract participants’ (N = 25, 13 females; mean age = 26.17, range: 20-33; mean arm length = 79.37 ± 6.07 cm) reachable limit by asking them to judge the reachability of the 2D visual stimuli adopted in Experiment I.

Procedures were identical to those of Experiment II, with the following exceptions. Participants could not see their hands, kept on their laps under the table. No tactile stimulation was delivered. They were asked to respond (reachable or not reachable) as fast as possible by pressing the left or the right foot pedal (order counterbalanced across subjects). A single block of 160 randomized trials was administered, including 16 visual (V) stimulations per position.

We computed the percentage of ‘reachable’ answers per position and fitted them to sigmoidal and normal curves, as in Experiment II. Through a paired t-test, we compared the root mean square error (RMSE) resulting from the fitting of a sigmoidal and a normal curve (both with 2 parameters) to these judgements. To test for possible differences in the temporal domain between the judgements of reachable and non-reachable positions, we analyzed participants’ RTs (Bourgeois & Coello, 2012) for each of the 10 positions by subjecting them to repeated measures ANOVA with Position (V-P1 to V-P10) as a within-subject factor.

Accuracy of the performance was globally high (> 99%). Comparing the RMSE resulting from the fitting of a sigmoidal and a normal curve to the judgements we obtained a significant difference (t_(24)_ = - 4.269, p < 0.001): residuals were smaller for the sigmoidal curve, indicating it provided a better fit for the data. When assessing the estimated coefficients of this curve, we obtained two parameters: the PSE (mean ± SE = 6.23 ± 0.15) and the slope (mean ± SE = −3.22 ± 0.55; see Figure S5).

As for Experiment 2, the percentages of “reachable” judgments were subjected to an ANOVA with Position (from V-P1 to V-P10) as within-subject factor. We observed a significant effect (F_(2.1,48.3)_ = 275.59, p < .001, η^2^_p_ = 0.92) that we further investigated though Tukey-corrected post-hoc. Again, the first five visual positions (not statistically different between themselves, all p_s_ > 0.05) reported significantly higher percentages of “reachable” responses compared to the last five (all p_s_ < 0.001). These results are in line with the objective limit of reachability of our participants (V-P6) and are represented in Figure S6.

The repeated measures ANOVA on RTs with Position (V-P1 to V-P10) as a factor did not report a significant effect: F_(1.25, 28.7)_ = 2.13, p = 0.152. Moreover, no differences between the RTs of judgement of reachable and unreachable positions emerged, preventing this variable from discriminating between stimuli within and beyond reach.

### Experiment S3

A main difference between Experiments I, II and S2 was the vision of the hand. Therefore, we ran Experiment S3 (N = 24, 14 females; mean age = 23.75, range: 18-38; mean arm length = 79.28 ± 5.10 cm) with the aim of introducing into our reaching task the same modulation of the hand we adopted in Experiments I and II. Previous studies adopting reachability judgement tasks to evaluate PPS extensions, in fact, required participants to place the hand in a visible position (Bourgeois & Coello, 2012). Therefore, we ran Experiment S3 identical to Experiment S2, but having the hand visible and in the same positions as in Experiment I. RTs and percentage of answers were computed the same way, but the close and distant hand conditions were run in separate blocks (160 randomized trials per block with the order counterbalanced).

As above in Experiment S2, sigmoidal and normal curves (both with 2 parameters) were separately fitted on the percentages of judgements, as a function of hand position. RTs were subjected to *Hand* (close vs. distant) * *Position* (V-P1 to V-P10) within-subject ANOVA.

Accuracy was high (>99%) and no differences between conditions emerged. In this case, the normal curve did not converge for 3 participants in the distant hand condition and for 2 participants in the close hand condition, whereas the sigmoidal curve provided a convergent fitting for all participants for both hand positions. Paired t-tests on the RMSE revealed a significant difference between the sigmoidal and the normal curve fitting, both for the distant (t_(20)_ = −5.32, p < 0.001) and for the close hand (t_(21)_ = −6.09, p < 0.001) positions: in particular, the sigmoidal curve reported the best fitting, as shown by smaller residuals, independent of the position of the hand. We then calculated the estimated coefficients of the sigmoid for each hand position, obtaining the PSE and the slope of the curve. Through a paired t-test, we compared the PSE for the distant (mean ± SE = 6.36 ± 0.15) and close (mean ± SE = 6.13 ± 0.22) positions; the difference was not significant (t_(23)_ = 1.446, p = 0.162), showing that the subjective judgement of the reachable boundary did not change depending on the hand location. We obtained the same result comparing the slope of the curves: t_(23)_ = −1.770, p = 0.090 (distant hand mean ± SE = −3.71 ± 0.65, close hand mean ± SE = −2.40 ± 0.44, see Figure S4). To study the contribution of the vision of the hand, we compared through an unpaired t-test the PSE and the slope values obtained in Experiment S2 (right hand not visible, placed on the right leg) with those obtained with the close hand in Experiment S3. The differences were not significant, both for the PSE (t_(40.24)_ = −0.405, p = 0.688) and for the slope (t_(45.39)_ = 1.17, p = 0.250).

**Figure S5:**
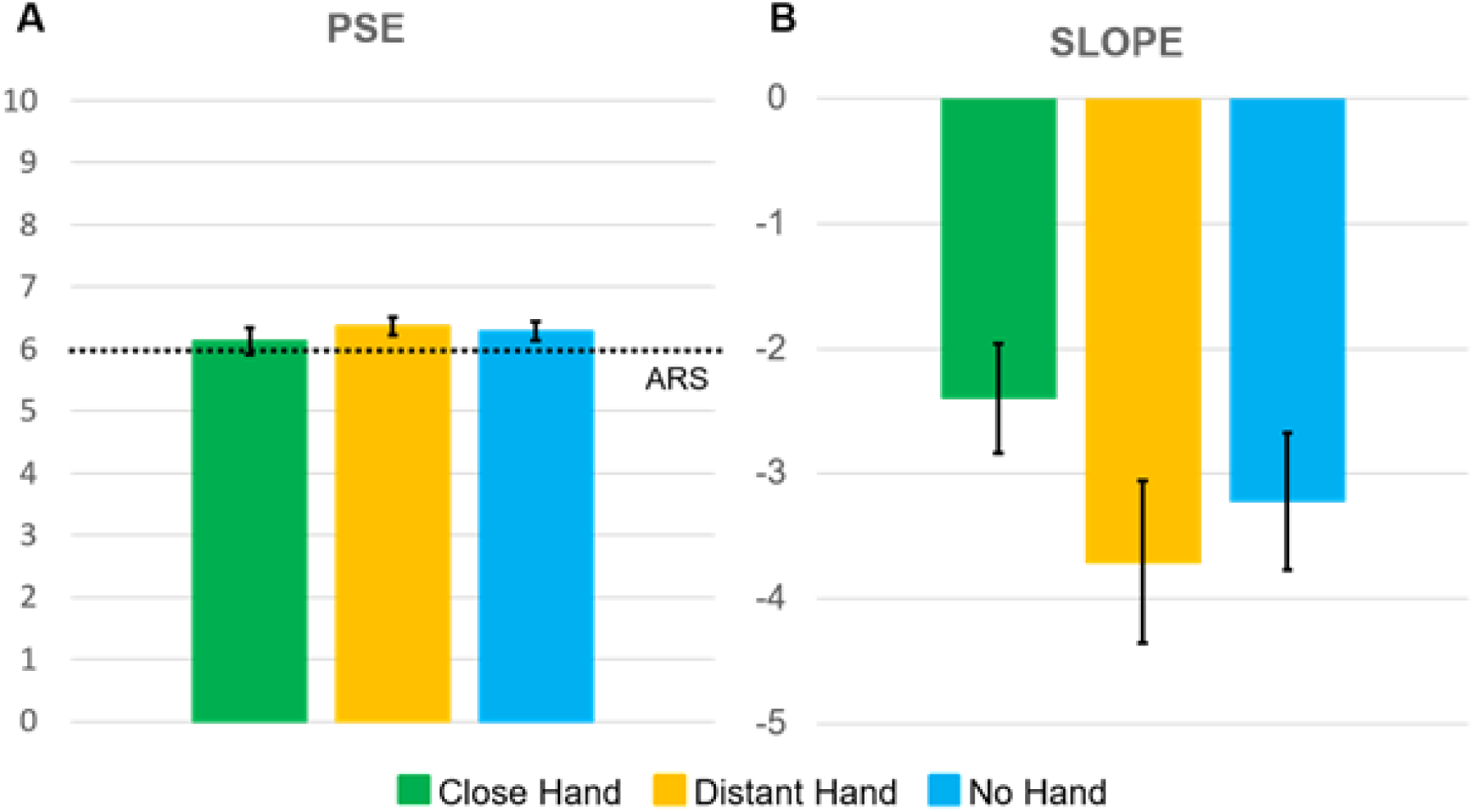
Subjective limit of reachability. a) PSE and b) slope values obtained in Experiment S2 (No Hand) and S3 (Distant or Close Hand). Error bars represent the standard error of the measure.

As done for Experiments II and S2, we performed a within-subject ANOVA 2 (Hand, Close vs Far) * 10 (Visual Position) on the percentages of “reachable” judgments issued by our participants. We observed a significant effect of Position (F_(2.06,47.33)_ = 268.62, p < .001, η^2^_p_ = 0.92) and a significant Hand * Position interaction (F_(4.14,95.14)_ = 3.43, p = .011, η^2^_p_ = 0.13). This interaction is significant due to a higher percentage of reachable responses in V-P5 and V-P6 (all p_s_ < 0.05, Bonferroni corrected) when the hand is distant. This could underline the fact that it is easier for participants to discriminate the reachability of a stimulation in border positions when the hand is close to that limit. These results are reported in Figure S6.

**Figure S6.**
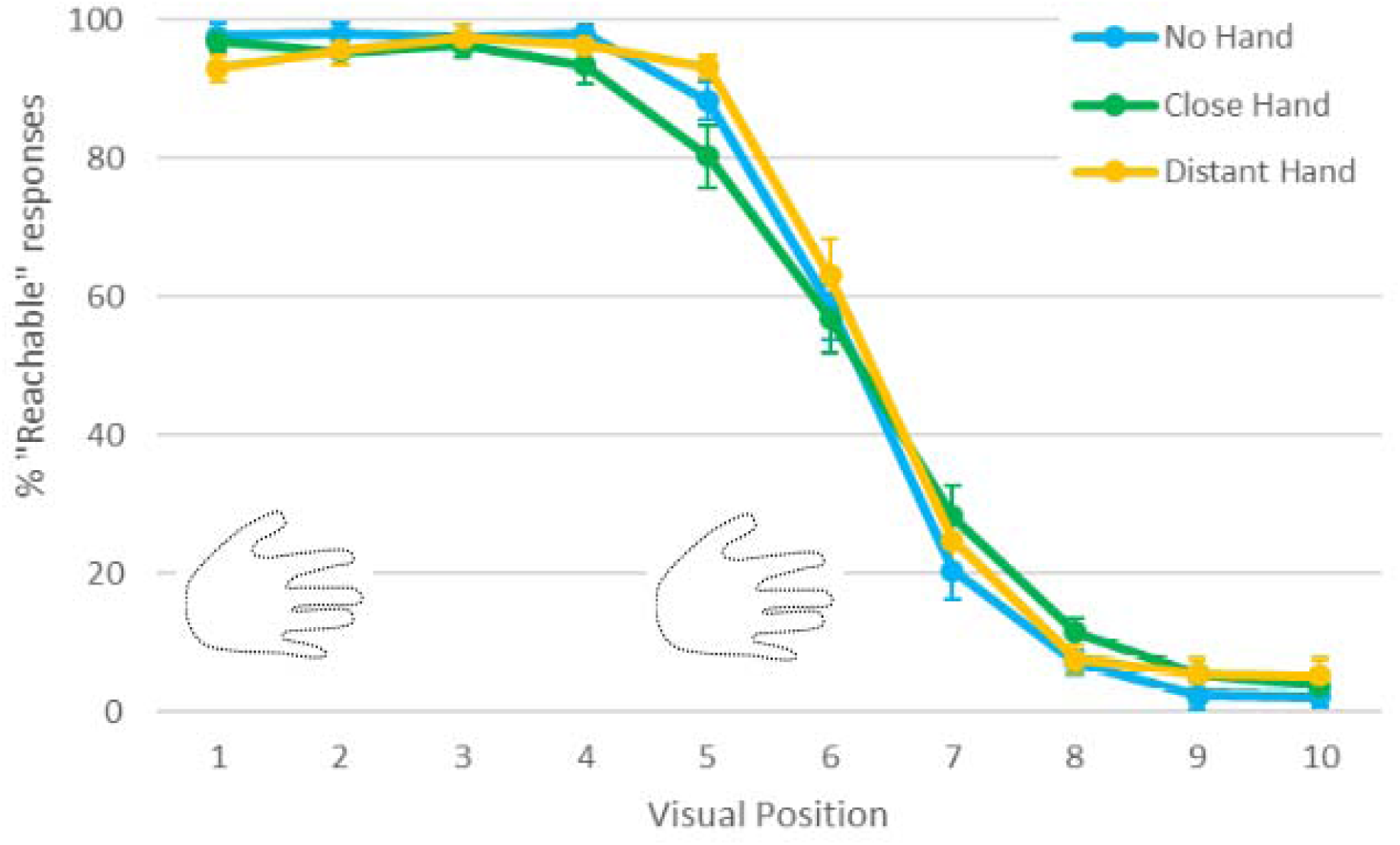
Percentages of reachability judgments along the 10 positions of visual stimulation for the close (green, Experiment S3), distant (yellow, Experiment S3) and no hand conditions (light blue, Experiment S2). The 6^th^ visual position represents the objective reachable limit. Error bars represent the standard error of the mean.

RTs were subjected to *Hand* (close vs. distant) * *Position* (V-P1 to V-P10) within-subject ANOVA. None of the main effects was significant: Hand (F_(1,23)_ = 0.67, p = 0.423) or Position (F_(1.40,32.27)_ = 1.23, p = 0.293). Similarly, the Hand*Position interaction was not significant (F_(4.75,109.16)_ = 0.68, p = 0.629). These results highlight the fact that reachability judgements over visual stimuli are not modulated by the position of the hand (Experiment S3) or by its visibility (comparing Experiment S2 with Experiment S3). This corroborates our hypothesis regarding the difference between PPS and ARS: contrary to what was observed in Experiment I, the position of the hand does not modulate the judgement speed.

